# Phagocyte-expressed glycosaminoglycans promote capture of alphaviruses from the blood circulation in a host species-specific manner

**DOI:** 10.1101/2023.08.09.552690

**Authors:** Stephanie E. Ander, M. Guston Parks, Bennett J. Davenport, Frances S. Li, Angela Bosco-Lauth, Kathryn S. Carpentier, Chengqun Sun, Cormac J. Lucas, William B. Klimstra, Gregory D. Ebel, Thomas E. Morrison

## Abstract

The magnitude and duration of vertebrate viremia are critical determinants of arbovirus transmission, geographic spread, and disease severity—yet, mechanisms determining arbovirus viremia levels are poorly defined. Previous studies have drawn associations between in vitro virion-glycosaminoglycan (GAG) interactions and in vivo clearance kinetics of virions from blood circulation. From these observations, it is commonly hypothesized that GAG-binding virions are rapidly removed from circulation due to ubiquitous expression of GAGs by vascular endothelial cells, thereby limiting viremia. Using an in vivo model for viremia, we compared the vascular clearance of low and enhanced GAG-binding viral variants of chikungunya (CHIKV), eastern-(EEEV), and Venezuelan-(VEEV) equine encephalitis viruses. We find GAG-binding virions are more quickly removed from circulation than their non-GAG-binding variant; however individual clearance kinetics vary between GAG-binding viruses, from swift (VEEV) to slow removal from circulation (EEEV). Remarkably, we find phagocytes are required for efficient vascular clearance of some enhanced GAG-binding virions. Moreover, transient depletion of vascular heparan sulfate (HS) impedes vascular clearance of only some GAG-binding viral variants and in a phagocyte-dependent manner, implying phagocytes can mediate vascular GAG-virion interactions. Finally, in direct contrast to mice, we find enhanced GAG-binding EEEV is resistant to vascular clearance in avian hosts, suggesting the existence of species-specificity in virion-GAG interactions. In summary, these data support a role for GAG-mediated clearance of some viral particles from the blood circulation, illuminate the potential of blood-contacting phagocytes as a site for GAG-virion binding, and suggest a role for species-specific GAG structures in arbovirus ecology.

**Significance Statement:** Previously, evidence of arbovirus-GAG interactions in vivo has been limited to associations between viral residues shown to promote enhanced GAG-binding phenotypes in vitro and in vivo phenotypes of viral dissemination and pathogenesis. By directly manipulating host GAG expression, we identified virion-GAG interactions in vivo and discovered a role for phagocyte-expressed GAGs in viral vascular clearance. Moreover, we observe species-specific differences in viral vascular clearance of enhanced GAG-binding virions between murine and avian hosts. These data suggest species-specific variation in GAG structure is a mechanism to distinguish amplifying from dead-end hosts for arbovirus transmission.

## Introduction

Viremia is an essential component for the lifecycle of mosquito-transmitted viruses. Maintenance of these viruses in nature requires infected vertebrate hosts to develop and maintain a viral titer in the blood sufficient for transmission to the mosquito host during a blood meal (1). As such, vertebrate species can be classified as either an amplifying or non-amplifying, “dead-end” host for a specific arbovirus depending on the magnitude and duration of viremia produced. Yet, the viral and host mechanisms that dictate the magnitude of vertebrate viremia, thereby distinguishing amplifying from non-amplifying hosts, have remained open questions.

Virion interactions with host glycosaminoglycans (GAGs) in the vasculature is one mechanism proposed to control arbovirus viremia (2–4). GAGs are long, linear polysaccharide chains and consist of five types defined by their repeating disaccharide units: heparan sulfate (HS), chondroitin sulfate (CS), dermatan sulfate (DS), keratan sulfate (KS), and hyaluronan (HA). As with other GAG-binding proteins, virion-GAG binding is attributed to electrostatic interactions of positively charged virion protein residues with the sulfated, negative-charged polysaccharide chains of GAGs (5). In vitro, many different viruses utilize GAGs as attachment factors, particularly HS (5–7), and genome-wide screens with arboviruses have highlighted a requirement of GAG biosynthetic pathways for chikungunya virus (CHIKV) (8), dengue virus (9), and Rift Valley fever virus (10) infections. In animal models of viremia, cell culture-adapted, enhanced GAG-binding arboviruses are more efficiently removed from the circulation than their non-enhanced GAG-binding (and non-cell culture adapted) progenitors (2–4, 11).

Based on these observations, it is commonly hypothesized that GAG-binding virions are rapidly removed from circulation due to ubiquitous expression of GAGs in the vasculature and thus limiting viremia and viral dissemination. However, this model of GAG-mediated viremic control neglects to account for the heterogeneity of GAG-binding observed amongst naturally circulating arbovirus isolates. For example, some natural and low-passaged isolates of Venezuelan equine encephalitis virus (VEEV), CHIKV, dengue virus, and Rift Valley fever virus display affinity for GAGs to various degrees (10, 12–16). Meanwhile, there is high conservation of GAG-interacting residues in the viral E2 glycoprotein amongst naturally circulating isolates of North American eastern equine encephalitis virus (EEEV) (17–19). These examples highlight a paradox with the current model: GAGs expressed on the host endothelium are proposed to restrict viremia development of GAG-binding arboviruses, yet GAG-binding viral isolates are maintained in natural circulation.

To investigate a role for GAGs in virus-host interactions in the vasculature, we employed a reductionist model of viremia. As described previously for this model (20), a defined number of viral particles (measured by RNase-resistant viral genomes) is directly inoculated into the bloodstream to simulate a viremic state, and the amount of viral particles remaining in circulation at various times post-inoculation is quantified. By directly inoculating virus into the bloodstream and analyzing samples before progeny virions are produced, this approach avoids interpretative complications that may arise both from differences in dissemination following a subcutaneous inoculation as well as de novo viral replication. Instead, the reductionist model of viremia provides a robust system to directly assess in vivo virus-host interactions in the vasculature. We previously applied this model to elucidate novel virus-host interactions of the arthritogenic alphaviruses CHIKV, Ross River virus, and o’nyong ’nyong virus with liver Kupffer cells mediated by the scavenger receptor macrophage receptor with collagenous structure (MARCO) (20).

Using this model for viremia, we compared the clearance of low and enhanced GAG-binding viral variants of CHIKV, EEEV, and VEEV. As expected, we found enhanced GAG-binding virions are more efficiently removed from the circulation than their non-enhanced GAG-binding variant. However, individual clearance kinetics vary between enhanced GAG-binding viruses, from swift (VEEV) to slower (EEEV) removal from circulation. Remarkably, we found phagocytic cells are required for the efficient clearance of enhanced GAG-binding virions, suggesting that unique, phagocyte-specific GAGs promote virion clearance. Furthermore, transient depletion of vascular HS specifically impedes the clearance of enhanced GAG-binding CHIKV and EEEV variants in a phagocyte-dependent manner. Surprisingly, the vascular clearance of enhanced GAG-binding VEEV particles is unaffected by vascular HS depletion, suggesting distinct virion-GAG interactions. Finally, extending the reductionist viremia model to an avian species (chicken), we observed HS-interacting EEEV particles are resistant to vascular clearance—in stark contrast to the phenotype observed in mice and suggestive of species-specific vascular GAG structures. In summary, this report provides striking evidence that heretofore putative arbovirus-GAG interactions do occur in vivo. Moreover, we demonstrate (i) viral specificity with select classes of GAGs in the vasculature, (ii) a distinct preference for phagocyte-expressed HS in the removal of CHIKV and EEEV particles from the circulation, and (iii) species-specific differences in the interactions of arboviruses with vascular GAGs.

## Results

To investigate the role of arbovirus-GAG interactions in clearance of viral particles from the bloodstream, we assembled a panel of isogenic alphaviruses with known characteristics or mutations that either confer or disrupt enhanced GAG-binding in vitro (**Table 1**). To permit non-select agent, mechanistic study of the interactions of EEEV and VEEV particles with host factors, we used Sindbis virus (SINV) chimeras encoding the structural open reading frame of EEEV (FL93) or VEEV (TrD or INH9813). As such, resultant virions generated from the chimera design appear structurally similar to wildtype (WT) EEEV or VEEV but are attenuated in animal models of viral pathogenesis (21, 22).

**Table 1.** Alphaviruses used in this study.

| **Virus** | **Abbreviation** | **Variant** | **Enhanced GAG-binding Residues** | **Ref.** |
| --- | --- | --- | --- | --- |
| Chikungunya virus | CHIKV | AF15561 (wt) | none |  |
|  |  | 181/25 | E2 glycoprotein G82R mutation | (24) |
| Venezuelan equine encephalitis virus | VEEV-IAB | TrD (wt) | none |  |
|  |  | TC-83 | E2 glycoprotein T120R mutation | (3, 64) |
|  |  | TrD<sup>T120R</sup> | E2 glycoprotein T120R mutation | (3, 64) |
|  | VEEV-IC | INH9813 (wt) | none |  |
|  |  | INH9813<sup>E3K</sup> | E2 glycoprotein E3K mutation | (25) |
| Eastern equine encephalitis virus | EEEV | FL93 (wt) | E2 glycoprotein K71, K74, K77 | (17) |
|  |  | K71-74-77A | none |  |

### Enhanced GAG-binding virions are efficiently removed from the circulation in a reductionist model of viremia

We assessed the ability of enhanced GAG-binding virions to be cleared from the circulation in comparison with their non-enhanced GAG-binding variants (**Figure 1**). In comparison with WT CHIKV AF15561, its derivative 181/25 is known to exhibit enhanced GAG-interactions conferred by a single point mutation in the E2 glycoprotein, G82R (23, 24). We found that both viruses are removed from the circulation within 1 hour post inoculation (hpi) with the enhanced GAG-binding CHIKV 181/25 exhibiting an accelerated initial clearance phase at 10 minutes pi (mpi) (**Figure 1A**). In contrast, the recognition and removal of encephalitic alphaviruses from circulation is entirely dependent on the presence of known enhanced GAG-binding residues (**Figures 1B-1D**). In experiments with VEEV subtypes IAB and IC, we observed the WT, virulent strains (TrD and INH9813) persist throughout the observed timepoints at high, steady levels in the circulation (**Figures 1C, 1D**). Meanwhile, variants of VEEV with known enhanced GAG-binding properties in the E2 glycoprotein are efficiently removed from circulation (**Figures 1C, 1D, Supplemental Figure 1**). These enhanced GAG-binding VEEV variants are characterized by the E2 glycoprotein mutations T120R in TC-83 (a derivative of TrD) (3) and E3K in INH9813 (25). WT EEEV is intrinsically capable of enhanced GAG-binding in vitro (17, 18), and this enhanced GAG-binding phenotype can be abrogated by alanine substitution of E2 lysines 71, 74, and 77 (K71-74-77A) (17, 18). In our reductionist model of viremia, we observed the presence of this lysine triad is critical for clearance of WT EEEV particles from the circulation (**Figure 1B**). Notably, while all four enhanced GAG-binding variants displayed faster clearance rates than their unenhanced GAG-binding counterparts, the former’s vascular clearance kinetics varied greatly among the virus panel (**Figure 1**). For example, the VEEV-IC mutant INH9813 (E3K) exhibited swift initial vascular clearance (**Figure 1D**), while WT EEEV was removed from the circulation at a slower rate (**Figure 1B**). These data suggest these enhanced GAG-binding viruses may bind different GAGs in vivo.

**Figure 1.**
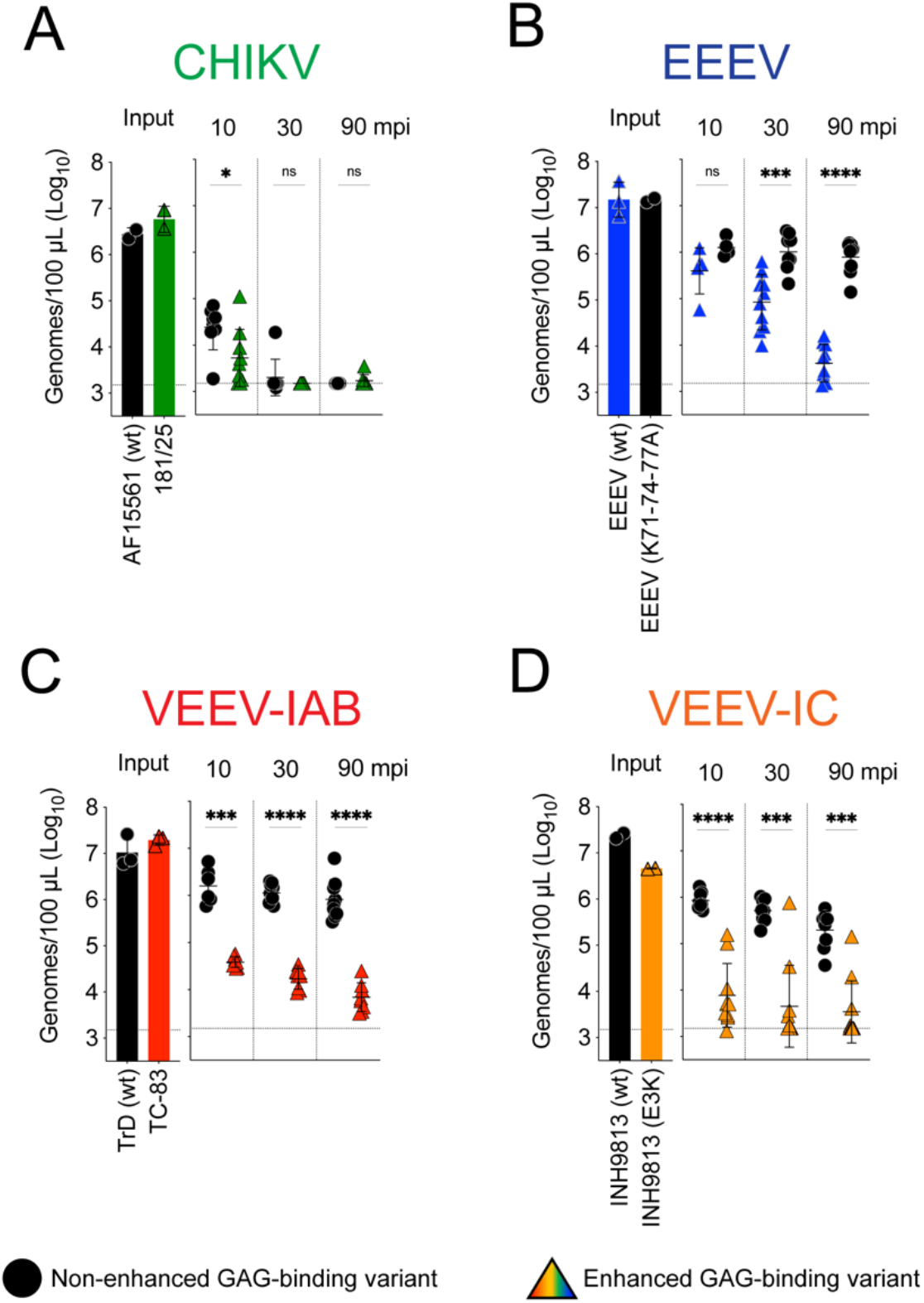
Virions with enhanced GAG-binding phenotypes in vitro display faster clearance kinetics from the circulation. Four-week old wildtype C57BL/6 mice were i.v. inoculated with viral particles (input) in a 100 μL volume. Serum was collected at 10, 30, and 90 minutes post-inoculation, and viral genomes were quantified by RT-qPCR. (**A**) Comparison of wildtype CHIKV (AF15561, n=8 mice per timepoint) with its enhanced GAG-binding derivative (181/25; n=9 mice per timepoint). (**B**) Comparison of wildtype, enhanced GAG-binding EEEV particles (SINV-EEEV chimera virus; n=10 mice per timepoint) with its loss of GAG-binding mutant (K71-74-77A SINV-EEEV chimera virus; n=9 mice per timepoint). (**C**) Comparison of wildtype VEEV-IAB particles (SINV-TrD chimera virus; n=6-9 mice per timepoint) with its enhanced GAG-binding derivative (TC-83; n=8 mice per timepoint). (**D**) Comparison of wildtype VEEV-IC particles (SINV-INH9813 chimera virus; n=7-8 mice per timepoint) with its enhanced GAG-binding mutant (SINV-INH9813 E2-E3K chimera virus; n=10 mice per timepoint). Data are combined from two independent experiments and displayed as mean ± standard deviation. Statistics are Mann-Whitney test; ****, p<0.0001; ***, p<0.001; *, p<0.05.

### Phagocytes contribute to the removal of enhanced GAG-binding virions from the circulation

We next assessed the role of phagocytes in mediating the clearance of enhanced GAG-binding virions from the circulation. To deplete phagocytes in direct contact with the blood, we intravenously (i.v.) administered clodronate-loaded liposomes (CLL) or PBS-loaded liposomes (PLL; control) at -2 days post inoculation (dpi) (26, 27). We then i.v. inoculated viral particles and assessed their clearance at 10 or 60 mpi, as determined by the vascular clearance kinetics shown in **Figure 1**. For a control, we included the WT, non-enhanced GAG-binder CHIKV AF15561 as we have previously established its clearance from the circulation is dependent on the presence of tissue-resident macrophages in the liver (20, 28). As shown in **Figures 2A** and **2C**, depletion of phagocytes in contact with the circulating blood impairs the clearance of the enhanced GAG-binding virions to varying degrees, and the greatest impact was observed on the vascular clearance of CHIKV 181/25 and WT EEEV.

**Figure 2.**
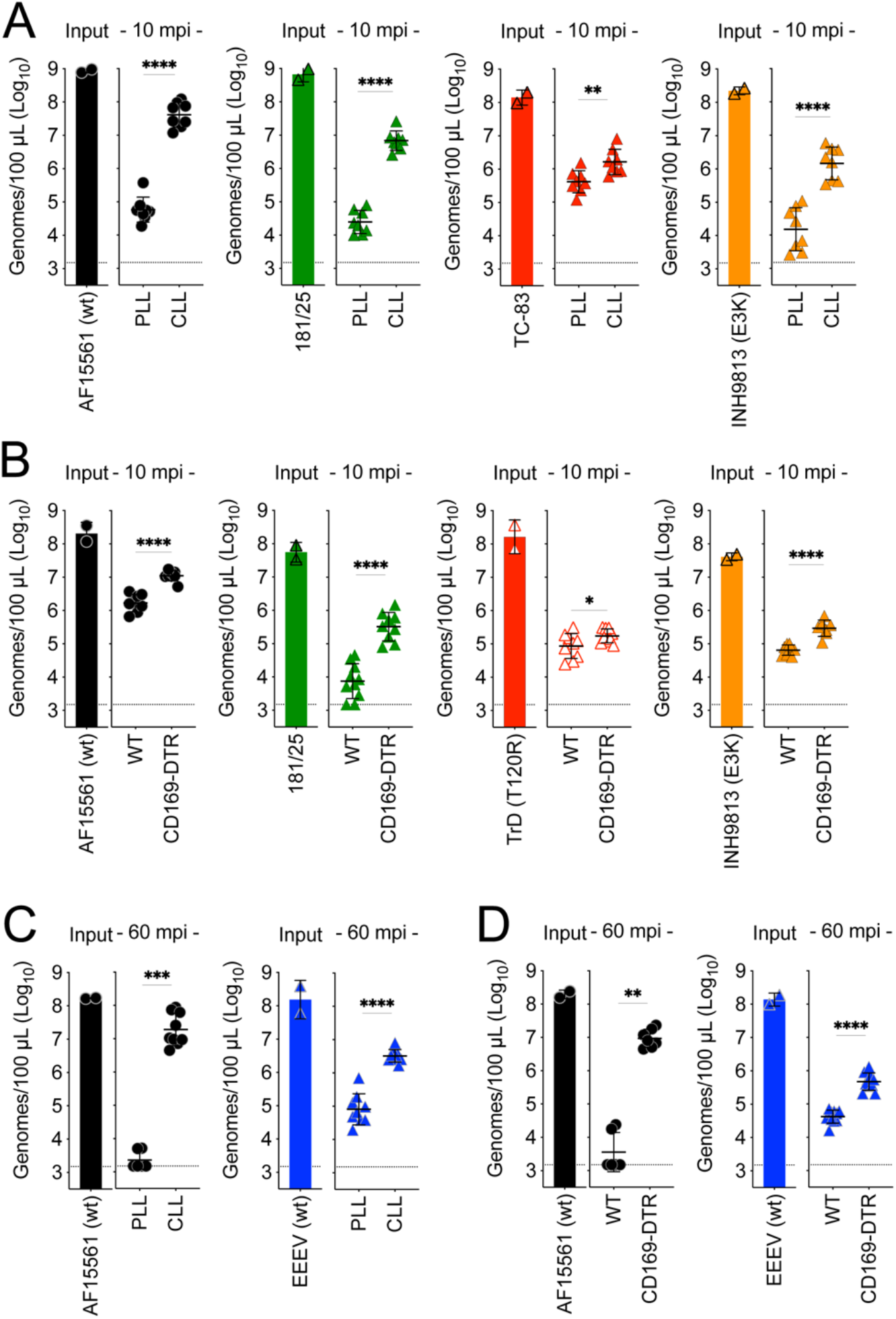
Depletion of phagocytes impairs clearance of enhanced GAG-binding virions from the circulation. (**A, C**) Two days prior to virus inoculation, wildtype (WT) mice were pre-treated with PBS-(PLL) or clodronate-loaded liposomes (CLL) i.v. (**B, D**) At 2 and 1 days prior to virus inoculation, WT and CD169-DTR mice were treated with DT i.p. (100 ng). (**A-D**) After the treatment period, mice were i.v. inoculated with viral particles (input) in a 100 μL volume. Serum was harvested at 10- or 60-minutes post-inoculation, and viral genomes were quantified by RT-qPCR. Data are combined from two independent experiments, with ≥ 3 mice/timepoint per experiment, and displayed as mean ± standard deviation. Statistics are unpaired T test or Mann-Whitney test; ****, p<0.0001; ***, p<0.001; **, p<0. 1; *, p<0.05.

We next investigated the role of tissue-resident macrophages by selectively depleting CD169^+^ cells using the CD169-DTR mouse model (29). Again, CHIKV AF15561 was used as a positive control (20). Intriguingly, we observed CD169^+^ cells to be significant contributors to the clearance of some, but not all, enhanced GAG-binding viruses (**Figures 2B, 2D**). Specifically, depletion of CD169^+^ cells significantly impedes the clearance of the enhanced GAG-binding variants CHIKV 181/25 and WT EEEV (**Figure 2D**). However, we observed minimal effect on the enhanced GAG-binding VEEV variants (**Figure 2B**). In summary, these data suggest that tissue-resident macrophages contribute to the viremic control of some, but not all, enhanced GAG-binding alphaviruses.

### The enhanced GAG-binding CHIKV variant 181/25 is removed from circulation by two independent pathways: GAG and MARCO interactions

Both WT CHIKV AF15561 and its enhanced GAG-binding derivative 181/25 are cleared efficiently from the circulation of WT mice (**Figure 1A**). Our previous study with CHIKV AF15561 identified both the host scavenger receptor MARCO (expressed by liver-resident macrophages) and the viral E2 glycoprotein residue K200 as necessary for viral clearance from murine circulation (20). However, in contrast to CHIKV AF15561, the enhanced GAG-binding CHIKV variant 181/25 is cleared from the circulation of MARCO^-/-^ mice, despite the conservation of the K200 residue (**Figure 3A**). Moreover, phagocytes are required for the clearance of CHIKV 181/25 in both WT and MARCO^-/-^ mice (**Figure 3B**). Based on these findings, we hypothesized CHIKV 181/25 is removed from the circulation by two independent, phagocyte-mediated pathways: (i) host MARCO interactions with viral residue K200 and (ii) host GAG interactions with viral residue R82.

**Figure 3.**
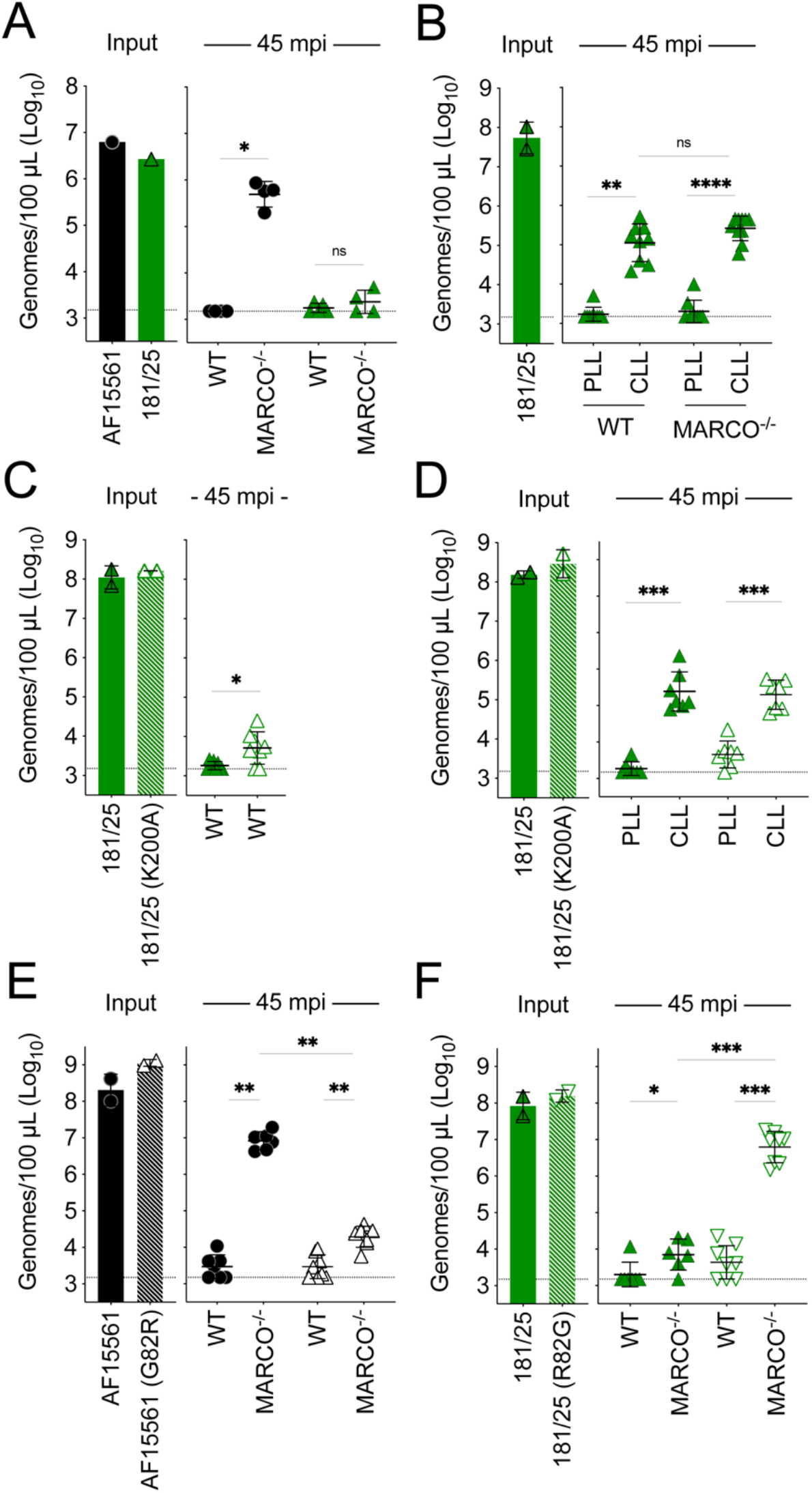
The enhanced GAG-binding mutation (E2-G82R) promotes MARCO-independent removal of CHIKV virions from the circulation. (**A-F**) Wildtype (WT) or MARCO^-/-^ C57BL/6 mice were i.v. inoculated with viral particles (input) in a 100 μL volume. Serum was collected at 45 minutes post-inoculation, and viral genomes were quantified by RT-qPCR. (**B, D**) Mice were pretreated with PBS-(PLL) or clodronate-loaded liposomes (CLL) i.v. at -2 dpi. Data are combined from 1-2 independent experiments, with 3-4 mice/timepoint per experiment, and displayed as mean ± standard deviation. Statistics are Mann-Whitney test; ****, p<0.0001; ***, p<0.001; **, p<0.01; *, p<0.05; ns, not significant.

To test this hypothesis and further establish a role for phagocytes in GAG-mediated clearance of virions from the circulation, we generated mutant viruses to selectively abolish viral interactions with MARCO or decrease GAG interactions. As our earlier work with CHIKV AF15561 (20) identified a stringent requirement for the lysine at position 200 of the CHIKV E2 glycoprotein for MARCO-mediated viremic control, we tested the effect of ablating this critical lysine residue on CHIKV 181/25 clearance. Congruent with the unaffected clearance of CHIKV 181/25 in MARCO^-/-^ mice (**Figure 3A**), the E2 K200A mutation in CHIKV 181/25 does not inhibit clearance of the virus from the circulation of WT mice (**Figure 3C**). Furthermore, we observed the vascular clearance of CHIKV 181/25 (K200A) is phagocyte-dependent, similar to 181/25 (**Figure 3D**). We next compared the effects of amino acid substitutions at viral E2 residue 82 in CHIKV AF15561 and 181/25 on viral vascular clearance in WT and MARCO^-/-^ mice. We observed clearance of CHIKV AF15561 from the circulation of MARCO^-/-^ mice is strictly dependent on the presence of the enhanced GAG-binding mutation E2-G82R (**Figure 3E**). Likewise, introduction of the E2-R82G mutation abrogates the clearance of CHIKV 181/25 in MARCO^-/-^ mice (**Figure 3F**). We thus concluded that viremic control of CHIKV 181/25 occurs by two separate host mechanisms, each sufficient to mediate clearance in the absence of the other: MARCO-dependent (requiring E2-K200) and MARCO-independent (requiring E2-R82). Moreover, phagocytes also play a significant role in the MARCO-independent, GAG-mediated removal of CHIKV 181/25 from the circulation.

### Depletion of vascular HS impairs the vascular clearance of enhanced GAG-binding CHIKV and WT EEEV, but not enhanced GAG-binding variants of VEEV

Previous in vitro studies with our alphavirus panel identified specific E2 glycoprotein residues that promote virion interactions with HS (**Table 1**) (3, 17, 18, 23–25). Therefore, to more definitively define the role of host HS in vivo, we assessed the effect of transient vascular HS-depletion on the vascular clearance of enhanced GAG-binding alphaviruses. At -1 hpi, we pretreated mice i.v. with a mixture of heparinase-I/heparinase-III to deplete vascular HS (30). We later confirmed treatment effectiveness by measuring freed syndecan-1 levels in the serum (**Supplemental Figure 3**) as i.v. heparinase injection causes syndecan-1 to become susceptible to extracellular matrix protease cleavage (31–33). Following heparinase treatment of WT mice, no effect was observed on the clearance of CHIKV 181/25 or AF15561 from the circulation (**Figure 4A, Supplemental Figure 2A**)—both of which retain the ability to interact with the host scavenger receptor MARCO via the viral E2 glycoprotein residue K200. In contrast, clearance of the enhanced GAG-binding CHIKV mutant 181/25 (K200A) is impaired by heparinase pre-treatment (**Figure 4B**). Moreover, heparinase treatment of MARCO^-/-^ mice impairs the vascular clearance of CHIKV 181/25 (**Figure 4C**)—supporting the hypothesis that the enhanced GAG-binding CHIKV variant 181/25 can be cleared from the circulation by both vascular GAGs and the scavenger receptor MARCO. Heparinase treatment also impedes clearance of enhanced GAG-binding, WT EEEV (**Figure 4D**), while no effect was observed on the non-enhanced GAG-binding EEEV mutant (K71-74-77A) (**Supplemental Figure 2B**). Notably, the effects of heparinase pre-treatment are virus-specific and cause no impediment to the vascular clearance of VEEV variants TrD (T120R) (**Figure 4E**) and INH9813 (E3K) (**Figure 4F**) nor their non-enhanced GAG-binding counterparts (**Supplemental Figure 2C, 2D**).

**Figure 4.**
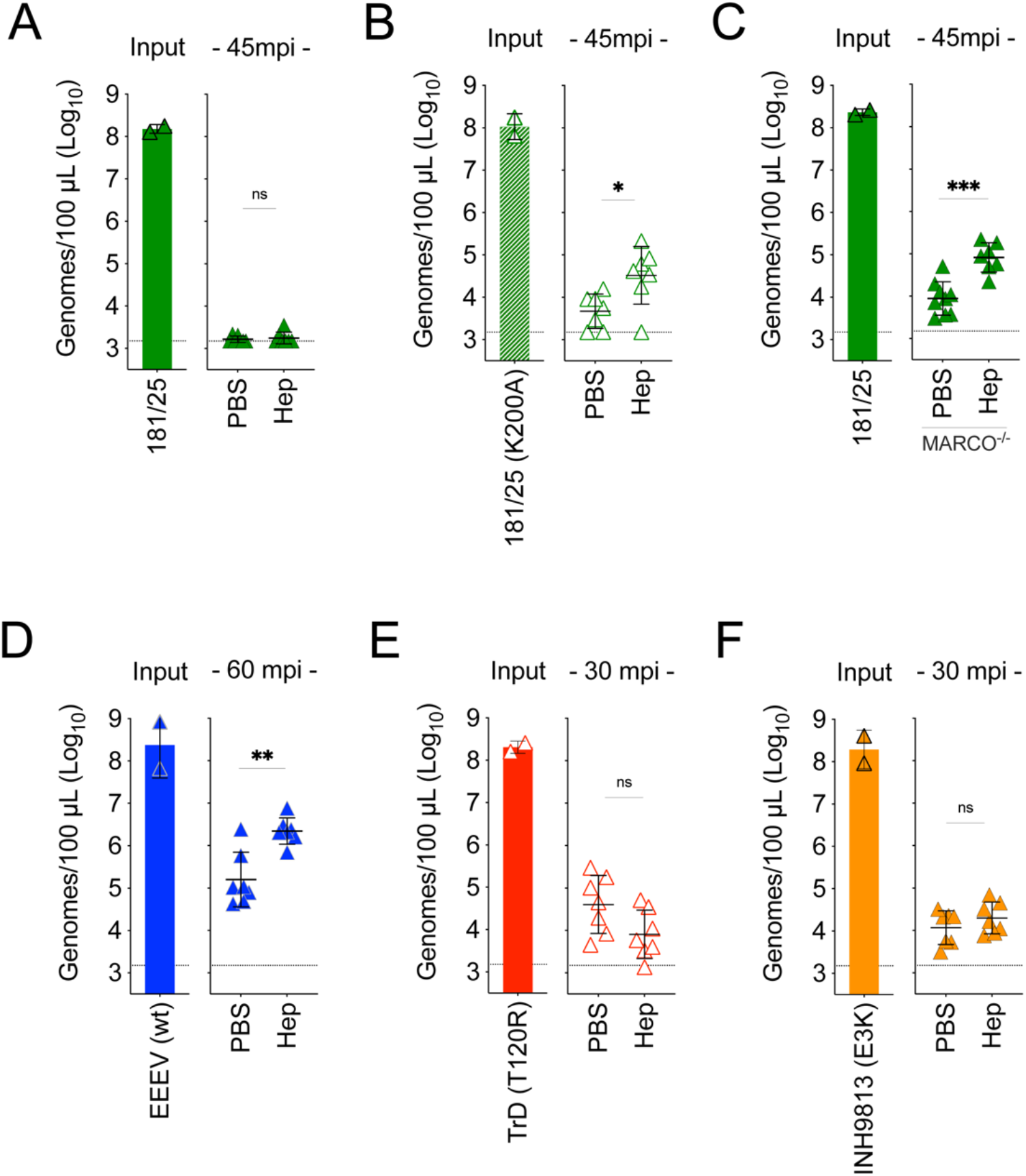
HS-deficiency in vivo impairs the removal of enhanced GAG-binding CHIKV and EEEV virions from the circulation but not enhanced GAG-binding VEEV variants. (**A-F**) Mice were pretreated i.v. with heparinase-I/III (Hep) or PBS; one hour later, they were i.v. inoculated with viral particles (input) in a 100 μL volume. Serum was collected at the indicated time post-inoculation, and viral genomes were quantified by RT-qPCR. (**A, B**) Vascular clearance of CHIKV variants in WT mice. (**C**) Vascular clearance of enhanced GAG-binding CHIKV 181/25 in MARCO^-/-^ mice. (**D**) Vascular clearance of wildtype EEEV (SINV-EEEV chimera) in WT mice. (**E-F**) Vascular clearance of enhanced GAG-binding VEEV variants (SINV-TrD and SINV-INH9813 chimeras) in WT mice. Data are combined from 2 independent experiments, with ≥ 3 mice/timepoint per experiment, and displayed as mean ± standard deviation. Statistics are Mann-Whitney test (**A**, **B**, **E**) and unpaired T test (**C**, **D**, **F**); ***, p<0.001; **, p<0.01; *, p<0.05; ns, not significant.

### Phagocyte-mediated removal of enhanced GAG-binding CHIKV and WT EEEV virions from the circulation is orchestrated by virion-HS interactions

Next, we assessed the role of phagocyte-specific HS expression by heparinase treatment of PLL- or CLL-treated mice. Following i.v. heparinase injection, phagocyte-intact mice (PLL group) display impaired clearance of the enhanced GAG-binding CHIKV mutant 181/25 (K200A), as expected (**Figure 5A**). However, i.v. heparinase treatment of phagocyte-depleted (CLL-treated) mice does not further impede viral clearance (**Figure 5A**). Likewise, similar experiments with WT EEEV also found the inhibitory effect of heparinase treatment is only detectable in the presence of phagocytes (**Figure 5B**). In total, these data suggest phagocyte expression of HS promotes the clearance of enhanced GAG-binding CHIKV and WT EEEV from the circulation (**Figure 6**).

**Figure 5.**
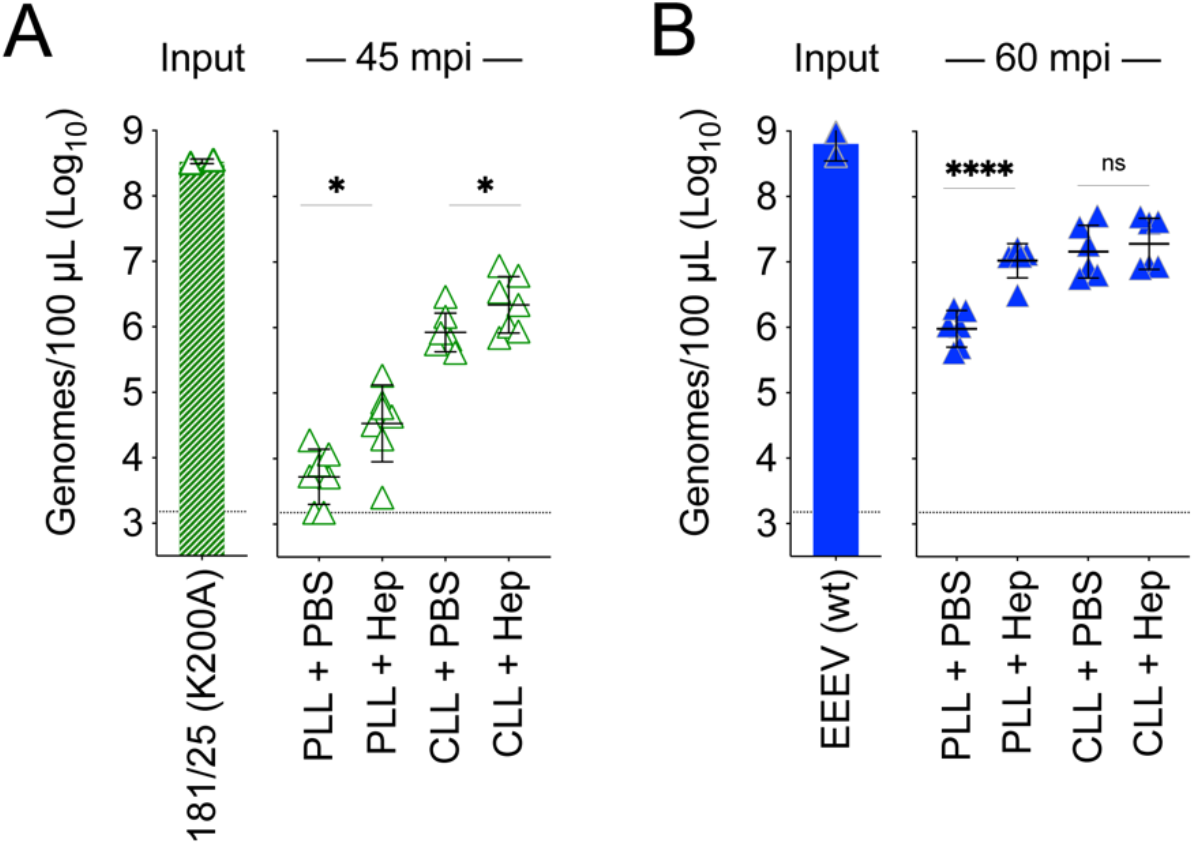
Phagocyte depletion eliminates the effect of vascular HS-deficiency on CHIKV and EEEV vascular clearance kinetics. At -2 dpi, WT mice were pretreated i.v. with PBS-(PLL) or clodronate-loaded liposomes (CLL); at -1 hpi, mice received a second pretreatment i.v. with heparinase-I/III (Hep) or PBS. All mice were then i.v. inoculated with viral particles (input) in a 100 μL volume. Serum was collected at the indicated time post-inoculation, and viral genomes were quantified by RT-qPCR. Data are combined from 2 independent experiments, with 3 mice/timepoint per experiment, and displayed as mean ± standard deviation. Statistics are Mann-Whitney test (**A**) or unpaired T test (**B**); ****, p<0.0001; *, p<0.05; ns, not significant.

**Figure 6.**
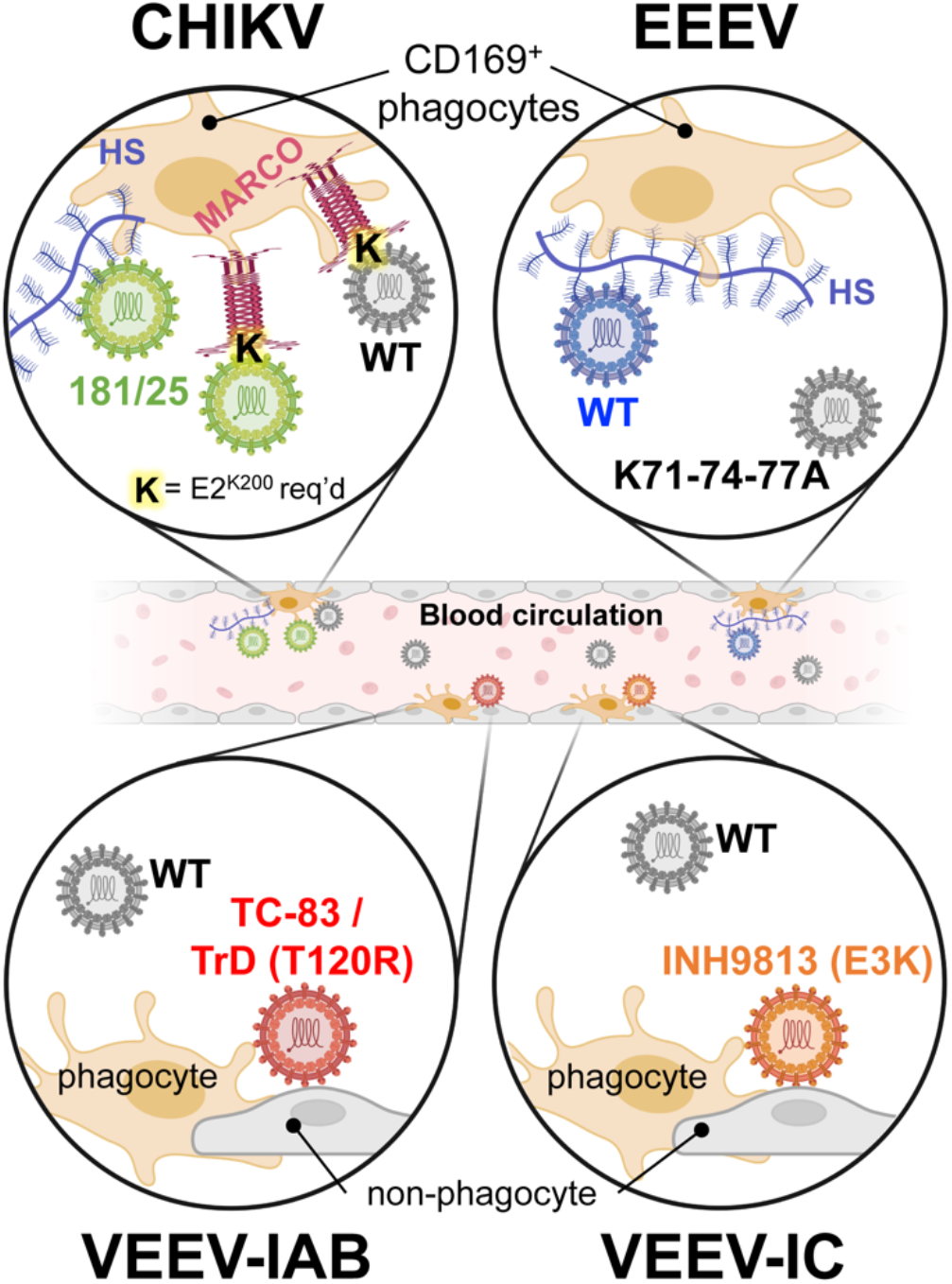
Model of the mechanisms by which enhanced-GAG binding virions are removed from murine circulation. CHIKV: enhanced GAG-binding variant 181/25 can be removed from murine circulation by tissue-resident macrophages by either HS- or MARCO-dependent mechanisms. EEEV: wildtype virus is captured by HS expressed on tissue-resident macrophages, while the non-enhanced GAG-binding K71-74-77A mutant evades vascular clearance. VEEV-IAB: the enhanced GAG-binding variants TC-83 and TrD (T120R) are removed from the circulation, and this is orchestrated by both phagocytes and non-phagocytic cells in a HS-independent manner. VEEV-IC: the enhanced GAG-binding variant INH9813 (E3K) is removed from the circulation by both phagocytes and non-phagocytic cells in a HS-independent manner.

### GAG-mediated viral vascular clearance is host species-dependent

The vascular clearance of GAG-binding arboviruses observed in this study and others (2–4, 11) raises the conundrum of how naturally circulating GAG-binding viruses, such as WT EEEV, maintain transmission cycles. Thus, we hypothesized that while GAGs are highly conserved throughout the animal kingdom (34, 35), species-specific variations in GAG structures (36–39) could produce differential viral vascular clearance phenotypes. To test this idea, we extended our reductionist viremia model to an avian system, as the natural enhanced GAG-binder WT EEEV is maintained in enzootic cycles between birds and mosquitoes (40). Previous studies have shown chicks develop a viremia capable of supporting mosquito transmission following subcutaneous WT EEEV inoculation (41–45). Thus, we i.v. inoculated WT EEEV and its non-GAG-binding mutant EEEV (K71-74-77A) into 6-9 day-old chicks and assessed viral vascular clearance at 1 and 3 hpi. Intriguingly, both the enhanced GAG-binding WT and the non-enhanced GAG-binding mutant EEEV virions are inefficiently cleared from chick circulation (**Figure 7A**). This inability of the avian vasculature to distinguish between the enhanced GAG-binding WT EEEV and its non-GAG-binding mutant is contrary to the distinct, opposing phenotypes observed in mice (**Figure 1B**). These observations of host-species dependent viral vascular clearance are especially striking considering EEEV ecology wherein avian species are natural amplifying hosts. We next assessed the vascular clearance of the VEEV variants, whose amplifying hosts are rodents and equines. Following i.v. inoculation into 6-9 day-old chicks, we observed both enhanced GAG-binding VEEV variants to be efficiently removed from the circulation while their non-enhanced GAG binding counterparts remained resistant to vascular clearance—mirroring the phenotypes we observed for these viruses in the mouse model (**Figure 7B and 7C**). Ultimately, these data suggest host species-specific HS structures in the vasculature may indeed be a contributing factor in defining amplifying from non-amplifying hosts (**Figure 7D**).

**Figure 7.**
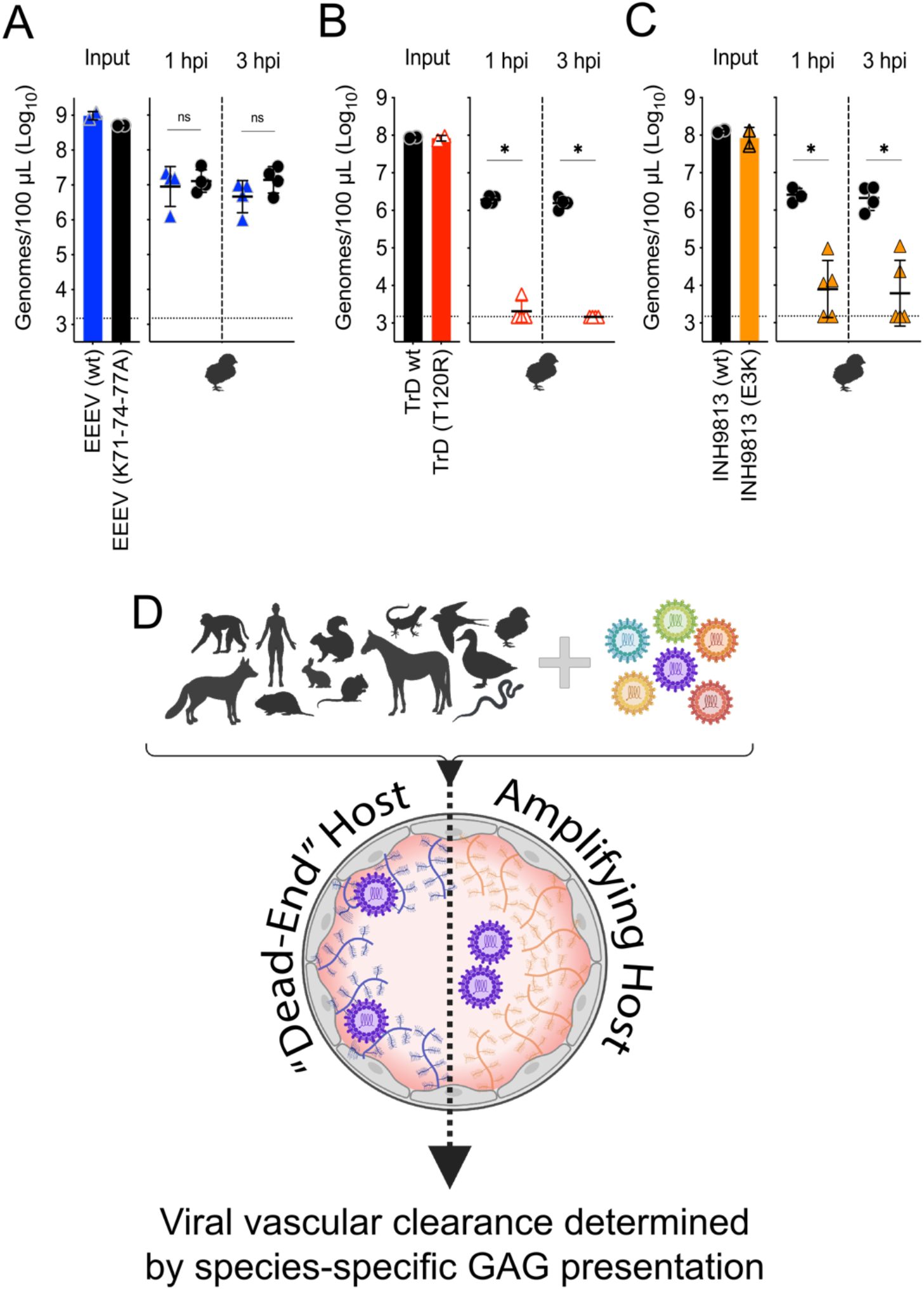
The vascular clearance phenotype of the natural enhanced GAG-binding alphavirus EEEV is species-specific. (**A-C**) 6-9 day-old chicks were i.v. inoculated with SINV-EEEV (**A**), SINV-VEEV-IAB (**B**), or SINV-VEEV-IC (**C**) viral particles (input) in a 100 μL volume. Serum was collected at the indicated times post-inoculation, and viral genomes were quantified by RT-qPCR. Data are combined from 2 independent experiments, with ≥2 chicks/virus per experiment, and displayed as mean ± standard deviation. Statistics are unpaired t test (**A**) and Mann-Whitney test (**C**, **D**); *, p<0.05 ns, not significant. (**D**) Model illustrating how variation in host-species GAG presentation may contribute to distinguishing “dead-end” hosts from amplifying hosts for individual arboviruses: arbovirus binding to GAGs only present in the vasculature of “dead-end” hosts (such as WT EEEV in the mouse) promotes capture of virions from the circulation, meanwhile the same arbovirus is able persist in the circulation of amplifying hosts (such as WT EEEV in the chick) due to species-specific differences in GAG presentation.

## Discussion

Previously, in vivo evidence of GAG interactions with a variety of arboviruses (including alphaviruses and flaviviruses) has been limited to associations between viral residues shown to promote enhanced GAG-binding phenotypes in vitro and in vivo phenotypes of viral dissemination and pathogenesis (2–4, 17, 23, 46–52). Meanwhile, the reciprocal experiments to alter host GAG expression have been impeded by observations that global genetic deletions of key GAG biosynthetic enzymes are embryonically lethal in mice (53, 54). This study supports the existence of arbovirus-GAG interactions in vivo through the direct manipulation of host GAG expression. Moreover, our studies also revealed a role for phagocyte-expressed GAGs and species-specific GAG structures in the clearance of arbovirus particles from the blood circulation (**Figures 6 and 7D**).

As expected, we observed virions with known enhanced GAG-binding phenotypes to be more efficiently removed from murine circulation than their non-enhanced GAG-binding variants (**Figure 1**). However, unlike previous studies (2–4, 11), our panel of enhanced GAG-binding viruses includes a representative of a naturally circulating enhanced GAG-binding virus, WT EEEV (**Figure 1B**). While also cleared from murine circulation, WT EEEV vascular clearance is slower than the other enhanced GAG-binding virions. This difference between a natural enhanced GAG-binding virion versus virions with enhanced GAG-binding conferred by cell culture adaptations suggests heterogeneity in the strength and/or specificity in virion-GAG interactions can influence the magnitude and duration of viremia.

Notably, we identified a distinct role for phagocytes in mediating the vascular clearance of enhanced GAG-binding virions. Following depletion of phagocytes in contact with the blood by clodronate-loaded liposomes, all enhanced GAG-binding virions were less efficiently removed from the circulation to varying degrees (**Figures 2 and 3**). These phenotypes were also corroborated in a mouse model of conditional and targeted ablation of CD169+ tissue resident macrophages (**Figure 2**). This data sits in contrast with the expectation that enhanced GAG-binding virions are trapped from the circulation by the ubiquitous expression of GAGs by endothelial cells (6). Instead, the efficiency of phagocyte-mediated vascular clearance of enhanced GAG-binding virions suggests that phagocytes express unique or a higher quantity of virus-binding GAGs than endothelial cells, or phagocytes may have faster turnover and replenishment of virus-trapping GAGs expressed on their surface.

To better define a role for host GAGs in alphaviral vascular clearance, we selectively depleted HS in the host vascular glycocalyx via i.v. pretreatments of mice with heparinase. In contrast with in vitro evidence that our panel of enhanced GAG-binding viruses bind to HS (3, 17, 24, 25), we found heparinase treatment did not impede the vascular clearance of all the enhanced GAG-binding alphavirus particles evaluated (**Figure 4**). This observation suggests in vivo variability in virion-GAG binding, and in vitro differential sensitivity to heparinase treatments has been previously described for the infectivity of EEEV, CHIKV, and SINV variants (18, 46, 49). Further investigation into the heparinase-sensitive viruses, CHIKV 181/25 (K200A) and WT EEEV, also found dual depletion of HS and phagocytes did not produce an additive effect on the level of vascular clearance inhibition (**Figure 5**)—suggesting that vascular clearance of CHIKV 181/25 (K200A) and WT EEEV is mediated by phagocyte-specific HS expression. Intriguingly, clearance of the enhanced GAG-binding VEEV mutants TrD (T120R) and INH9813 (E3K) is unaffected by vascular HS depletion (**Figure 4**), which suggests either insufficient HS depletion or a possible preference for other, non-HS GAGs in the context of the host vascular system.

We note a caveat of heparinase treatment, which is the subsequent susceptibility of some heparan sulfate proteoglycans to protease attack by sheddases within the extracellular matrix. For example, we utilized serum levels of the proteoglycan syndecan-1 as an indicator for heparinase activity. Syndecan-1 is presented at the cellular surface, but following heparinase treatment, it is shed into the extracellular milieu due to the unmasking of protease cleavage sites formerly shielded by the presence of its HS posttranslational modifications (31–33). It is therefore possible that enhanced GAG-binding CHIKV and WT EEEV may interact with the proteinaceous core of a heparinase-sensitive proteoglycan rather than HS itself. However, compelling in vitro data exists in the literature demonstrating HS binding by CHIKV 181/25 and WT EEEV using HS-coated agarose beads or cell lines deficient in GAG biosynthetic enzymes (17, 18, 23, 24).

Finally, the striking contrast of WT EEEV vascular clearance in a murine model to its vascular persistence in an avian model implies species-specificity in GAG structures may be a driver in determining which vertebrate species can participate in arboviral transmission (**Figure 7**). If so, this may explain the seeming paradox of the existence of naturally circulating arbovirus isolates with GAG-binding properties. From studies comparing the GAG composition of various tissues amongst vertebrate species, it is clear that there exists both tissue and species heterogeneity in the expression and structural features of HS and other GAGs (36–39). This heterogeneity can result from variations in polysaccharide chain length and disaccharide modifications (e.g., *N*-, 2-*O*-, 3-*O*-, and 6-*O*-sulfations found on HS molecules), as well as the specific pattern of these disaccharide modifications along the polymer (53, 55). This structural variation promotes a diversity of biological functions, wherein discrete regions of a GAG may present unique protein-binding sites (53, 55). Such specificity in HS sulfation for virus recognition has been characterized for SARS-CoV-2 and human cytomegalovirus (56, 57). As for why the enhanced GAG-binding variants of VEEV were efficiently cleared from the circulation of both murine and avian hosts, we hypothesize these cell-culture adapted viruses may either bind vascular GAGs common to both vertebrate species or exhibit more promiscuous GAG-binding overall.

Relatedly, it is intriguing to consider whether individual variation in GAG expression or structural features may be a determinant in the estimated 4% of human EEEV infections that develop into clinical disease (58). For example, the non-GAG binding EEEV mutant is associated with increased viremia and morbidity following subcutaneous inoculation of mice (17), presumably due to decreased virion-GAG interactions. As such, it might similarly be hypothesized that alterations in GAG structures due to genetic polymorphisms of GAG biosynthetic or modifying enzymes (59) or factors such as age, diet, or inflammation (60) could also influence viral disease severity in human infections.

In summary, this study highlights the existence of distinct mechanisms by which vertebrate hosts can exert viremic control over virions with enhanced GAG-binding properties. We observed both HS-dependent and -independent viral vascular clearance, suggesting a role for other classes of GAGs in mediating arboviral interactions in vivo. We also identified a role for phagocytes as a scaffold for HS-arbovirus interactions in the vasculature. Finally, our cross-species comparison of viral vascular clearance underscores the importance of investigating viral phenotypes in various contexts in order to achieve a more complete understanding of arbovirus biology and ecology.

## Materials and Methods

### Cells

BHK-21 cells (ATCC CCL10) were cultured at 37°C in α-Minimum Essential Medium (Gibco) supplemented with 10% FBS, 10% tryptose phosphate broth, and penicillin-streptomycin.

### Viruses

All viruses used in this study were generated from cDNA clones. Viruses used were CHIKV strains AF15561, 181/25, AF15561 E2-G82R, 181/25 E2-R82G, 181/25 E2-K200A; SINV-EEEV (FL93 strain) and SINV-EEEV E2-K71-74-77A; TC-83; SINV-VEEV-IAB (TrD strain) and SINV-VEEV-IAB E2-T120R; SINV-VEEV-IC (INH9813 strain) and SINV-VEEV-IC E2-E3K. CHIKV AF15561 E2-G82R and 181/25 E2-R82G have been previously described (23). CHIKV 181/25 E2-K200A was generated by site-directed mutagenesis of the parental 181/25 plasmid using the following primers: 5′-CAGACGGTGCGGTACGCGTGTAATTGCGGTGACTC-3′ and 5′-GAGTCACCGCAATTACACGCGTACCGCACCGTCTG-3′. SINV-EEEV was described previously (61) and was used to generate SINV-EEEV E2-K71-74-77A by restriction cloning using the E2-K71-77A full-length EEEV plasmid previously described. SINV-VEEV-IAB was previously described (61) and used to generate SINV-VEEV-IAB E2-T120R, also referred to as TrD (T120R), by site-directed mutagenesis with the following primers: 5′-GAGCAGGAGTGTCTGACGGAATCTTTCTTAAATTCC-3′ and 5′-GGAATTTAAGAAAGATTCCGTCAGACACTCCTGCTC-3′. The enhanced GAG-binding SINV-VEEV-IC mutant was generated based on the previously described full-length VEEV-IC recombinant virus (25) and constructed similarly as described for SINV-VEEV-IAB (61). The wildtype SINV-VEEV-IC sequence was based on the previously described full-length SINV-VEEV-IC (25) and generated by site-directed mutagenesis of the enhanced GAG-binding variant using primers 5′-GAAGAAAAAGGAGATCTACCGAGGAGCTGTTTAAGGAGTAT-3′ and 5′-ATACTCCTTAAACAGCTCCTCGGTAGATCTCCTTTTTCTTC-3′. To generate virus stocks, plasmids containing the viral cDNA were linearized and used as the template for SP6 RNA polymerase *in vitro* transcription (mMessage mMachine). The RNA products were then electroporated into BHK-21 cells, and clarified supernatant was harvested at 24-30 h post-electroporation and stored as aliquots at -80°C. Infectious virus titers were determined by plaque assay on BHK-21 cells (62). Viral genomes were quantified by RT-qPCR of RNA extracted from virus stocks, as described below.

### Mouse Experiments

All mouse studies were performed following the recommendations in the Guide for the Care and Use of Laboratory Animals of the National Institutes of Health. The protocols were approved by the Institutional Animal Care and Use Committee of the University of Colorado School of Medicine (Assurance Number A3269-01). Four-week old mice were used in all experiments. WT C57BL/6J mice were ordered from The Jackson Laboratory (strain #000664); CD169-DTR (29) and MARCO^-/-^ (63) C57BL/6 mice were bred in-house. All WT mice were age-matched males and were randomly assigned to experimental groups; prior studies did not identify sex-based differences. In-house bred mice of both sexes were used and randomly assigned to experimental groups to produce approximately equal sex distribution. All animal experiments were performed in ABSL2 or ABSL3 conditions as required. For phagocyte depletion, clodronate-loaded liposomes (ClodronateLiposomes.org) were administered i.v. at 10 µL/g on -2 dpi (20). PBS-loaded liposomes (ClodronateLiposomes.org) were likewise administered as controls. For targeted depletion of tissue-resident phagocytes, CD169-DTR and WT (control) mice received i.p. injections of DT (100 ng) on -2 and -1 dpi. To deplete heparan sulfate from the vascular glycocalyx, mice were i.v. injected at -1 hpi with 0.042 IU of recombinant heparinase I and III mixture, as described previously (30). Heparinase efficacy *in vivo* was verified by syndecan-1 ELISA (Abcam ab273165) of serum. For viral vascular clearance experiments, isoflurane vapors were used as anesthesia and mice were inoculated i.v. with 10^7^-10^9^ viral genomes diluted to a final 100 µL volume in PBS/1% FBS. At indicated timepoints, mice were humanely euthanized and blood was collected for serum isolation.

### Chick Experiments

Chick study protocols were approved by the Institutional Animal Care and Use Committee of Colorado State University (Assurance Number A3572-01). 6-9 day-old outbred chicks of indeterminant sex were used in vascular clearance experiments and obtained from Northern Colorado Feeder Supply. Chicks were i.v. inoculated with 10^8^-10^9^ viral genomes diluted to a final 100 µL volume in PBS/1% FBS. Blood was serially collected at 1 and 3 hpi for serum isolation.

### Viral titer calculations by RT-qPCR

Viral genomes were quantified from 20 µL of serum. Briefly, 20 µL of serum were placed into TRIzol, and RNA was extracted using the PureLink RNA Mini Kit (Life Technologies). SuperScript IV reverse transcriptase (Life Technologies) was used to generate cDNA from 10 µL of serum-derived RNA, and viral genomes were quantified by qPCR using the following primers and probes: CHIKV forward primer (5′-TTTGCGTGCCACTCTGG-3′), CHIKV reverse primer (5′-CGGGTCACCACAAAGTACAA-3′), and CHIKV internal TaqMan probe (5′-ACTTGCTTTGATCGCCTTGGTGAGA-3′); SINVnsp2 forward primer (5′-ACGCCGACCGCAACAGTGAG-3′), SINVnsp2 reverse primer (5′-TCTCGCTGCGGACACCCTGA-3′), and SINVnsp2 internal TaqMan probe (5′-ACGTAGTCACCGCTCTTGCCA-3′); TC-83 forward primer (5′-GCCTGTATGGGAAGCCTTCA-3′), TC-83 reverse primer (5′-TCTGTCACTTTGCAGCACAAGAAT-3′), and TC-83 internal TaqMan probe (5′-CCTCGCGGTGCATCGTAGCAGC-3′). As described previously, a standard curve was generated using in vitro synthesized RNA; this was used to determine the absolute quantity of viral RNA in experimental samples (62). The standard curve for quantification of TC-83 was generated by 10-fold serial dilutions of a plasmid encoding the TC-83 genome from 10^8^ to 10^0^ copies.

## Acknowledgments

This work was funded by Public Health Service Grants T32 AI007405 and F32 AI161866 to SEA, F30 AI160828 to FSL, F32 AI140567 to KSC, R01 AI095436 to WBK, R01 AI067380 to GDE, and R01 AI148144 to TEM. The funders had no role in study design, data collection and analysis, decision to publish, or preparation of the manuscript. We would also like to thank Kaori Oshima and Eric Schmidt (Massachusetts General Hospital) for the gift of the recombinant heparinase-I and -III used in this study. Diagrams were created with Biorender.com.

