## Supplemental Figures 1-3 for "Phagocyte-expressed glycosaminoglycans promote capture of alphaviruses from the blood circulation in a host species-specific manner"

*Thomas E. Morrison

**Author Contributions:** SEA, TEM, and GDE designed experiments; SEA, BJD, FSL, ABL, KSC, and GDE performed experiments; SEA and MGP processed samples; SEA, CS, CJL, and WBK generated/supplied reagents; SEA and TEM analyzed data; SEA wrote initial manuscript draft; SEA, ABL, WBK, GDE, and TEM edited manuscript.

**Keywords:** viremia, glycosaminoglycan, heparan sulfate, alphavirus, arbovirus.

**This PDF file includes:**

Supplemental Figures 1 to 3

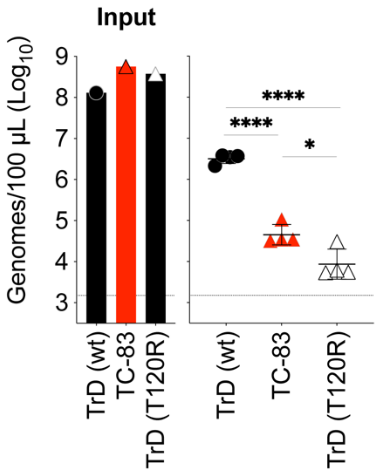


**Supplemental Figure 1. The E2-T120R enhanced GAG-binding mutation in TC-83 is sufficient to promote the clearance of TrD from the circulation.** The enhanced GAG-binding mutation in the E2 glycoprotein of VEEV TC-83 (T120R) was introduced into the SINV-TrD chimera. Mice were inoculated with a defined number of viral particles (genomes) in a 100 µL volume. Serum was collected at 90 minutes post-inoculation, and viral genomes were quantified by RT-qPCR. Data shown is from 1 experiment, with 4 mice/timepoint, and displayed as mean ± standard deviation. Statistics are unpaired T test; ****, $p<0.0001$; *, $p<0.05$.

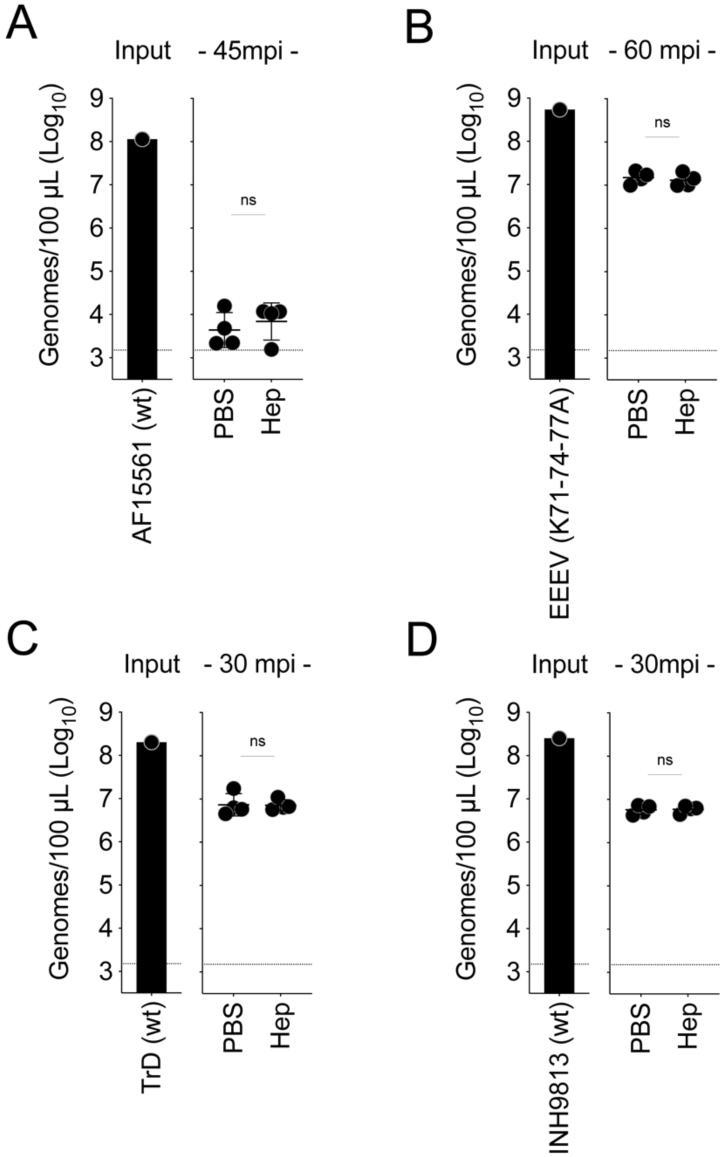


**Supplemental Figure 2. Transient depletion of HS does not alter vascular clearance phenotypes of non-enhanced GAG-binding virions.** (**A-D**) At -1 hpi, wildtype mice were i.v. injected with heparinase-I/III (Hep) or PBS to cleave heparan sulfate GAGs in contact with the blood. Mice were then inoculated with a defined number of viral particles (genomes) in a 100 µL volume. Serum was collected at indicated times post-inoculation, and viral genomes were quantified. Data shown is from 1 experiment, with 4 mice/group, and displayed as mean ± standard deviation. Statistics are Mann-Whitney test; ns, not significant.

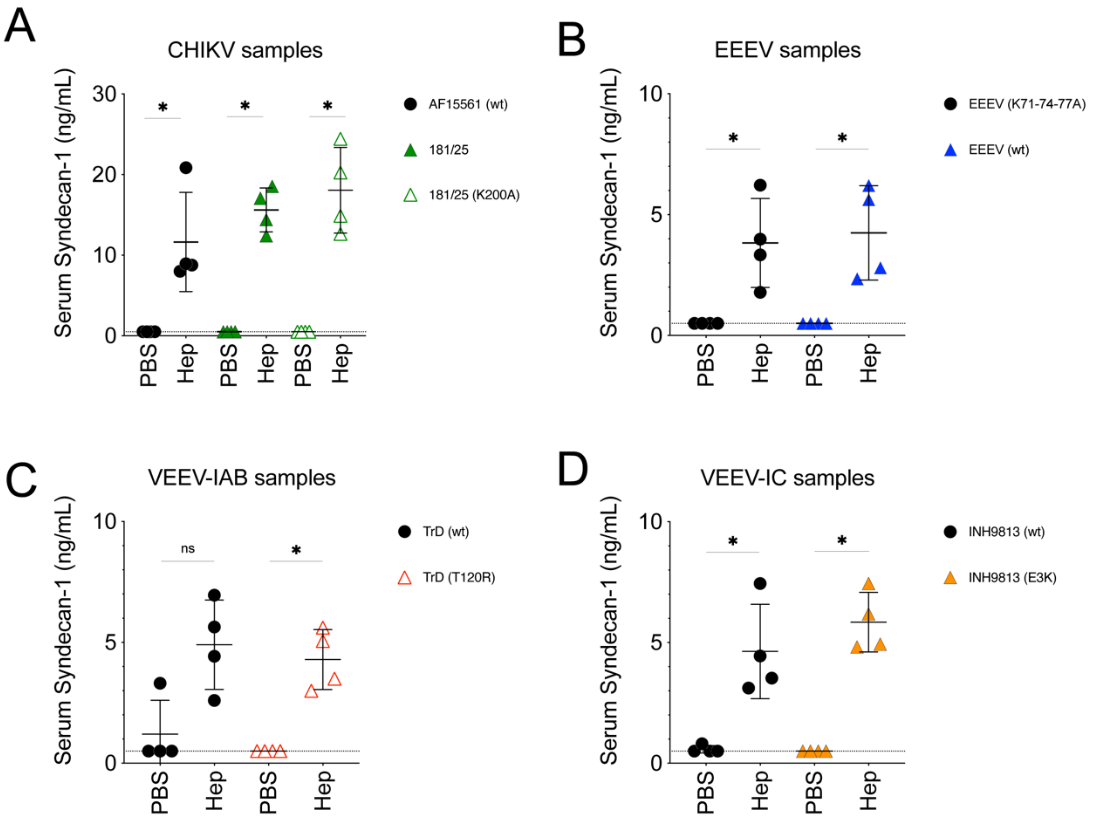


**Supplemental Figure 3. Treatment of mice with i.v. heparinase-I/III induces shedding of the proteoglycan syndecan-1, a read-out of enzyme effectiveness.** (**A-D**) At -1 hpi, wildtype mice were i.v. injected with heparinase-I/III (Hep) or PBS to cleave heparan sulfate GAGs in contact with the blood. Mice were then inoculated with a defined number of viral particles (genomes) in a 100 µL volume. Serum was collected at 30 (**C, D**), 45 (**A**), or 60 (**B**) minutes post-inoculation, and the syndecan-1 level in the serum was measured by ELISA. Heparinase treatment of cells removes HS modifications from the proteoglycan syndecan-1, thus exposing cleavage sites to proteases in the extracellular matrix. Data shown is from 1 experiment, with 4 mice/group, and displayed as mean ± standard deviation. Statistics are Mann-Whitney test; *, $p<0.05$; ns, not significant.
